# Periodic synchronization of dengue epidemics in Thailand: the roles played by temperature and immunity

**DOI:** 10.1101/2021.03.01.433325

**Authors:** Bernardo García-Carreras, Bingyi Yang, Mary K. Grabowski, Lawrence W. Sheppard, Angkana T. Huang, Henrik Salje, Hannah Eleanor Clapham, Sopon Iamsirithaworn, Pawinee Doung-Ngern, Justin Lessler, Derek A. T. Cummings

**Affiliations:** Department of Biology, University of Florida, Gainesville, FL, USA; Emerging Pathogens Institute, University of Florida, Gainesville, FL, USA; Department of Pathology, Johns Hopkins School of Medicine, Baltimore, MD, USA; Department of Ecology and Evolutionary Biology and Kansas Biological Survey, University of Kansas, Lawrence, KA, USA; Department of Virology, Armed Forces Research Institute of Medical Sciences, Thailand; Department of Genetics, University of Cambridge, Cambridge, UK; Saw Swee Hock School of Public Health, National University of Singapore, Singapore; Department of Disease Control, Ministry of Public Health, Nonthaburi, Thailand; Department of Epidemiology, Johns Hopkins Bloomberg School of Public Health, Baltimore, MD, USA

## Abstract

The spatial distribution of dengue and its vectors (spp. Aedes) may be the widest it has ever been, and projections suggest that climate change may allow the expansion to continue. However, the largest impacts of climate change on dengue might be in regions where the pathogen is already endemic. In these areas, the waxing and waning of immunity has a large impact on temporal dynamics of cases of dengue haemorrhagic fever. Here, we use 51 years of data across 72 provinces and characterise spatio-temporal patterns of dengue in Thailand, where dengue has caused almost 1.5 million cases over the last thirty years, and examine the roles played by temperature and dynamics of immunity in giving rise to those patterns. We find that timescales of multiannual oscillations in dengue vary in space and time and uncover an interesting spatial phenomenon: Thailand has experienced multiple, periodic synchronization events. We show that patterns in synchrony of dengue are consistent with those observed in temperature. Applying a temperature-driven dengue model, we explore how dynamics of immunity interact with temperature to produce the observed multiannual dynamics and patterns in synchrony. While multiannual oscillations are readily produced by immunity in absence of multiannual timescales in temperature, synchrony in temperature can synchronise dengue dynamics in different locations. However, at higher mean temperatures and lower seasonal variation, immune dynamics become more predominant, and dengue dynamics become more insensitive to multiannual fluctuations in temperature. These findings can help underpin predictions of disease patterns as global temperatures rise.

**Author summary:** 

## Introduction

The spatio-temporal dynamics of animal populations, including fluctuations and correlations in amplitudes and phases, has been an important area of research in ecology and the physical sciences for decades. Empirical observations of patterns in population dynamics have propelled theory forward leading to a better understanding of predator-prey interactions [1, 2], allee effects [3] and the interactions between deterministic and stochastic elements of systems [4, 5]. While a rich literature exists on synchronization of systems and their causes [6], there is little empirical evidence for periodic oscillations or fluctuations in spatial synchrony, particularly in epidemiology, where identification of long-term patterns in infectious disease dynamics could be critical for the success of health interventions. For instance, knowing when pathogen population levels are particularly low concurrently across a region presents improved opportunities for pathogen elimination [7]. Likewise, anticipation of global epidemics may assist in the structured allocation of resources, such as vaccines, across space and time. While regular spatial synchrony in other wildlife species has been described (e.g., [8–12]), how the degree of synchrony may vary over time has received less attention [13–15]. Here, we describe regular periodic synchronization in the dynamics of dengue haemorrhagic fever (DHF) in Thailand over a 51-year interval.

Dengue virus (DENV) is a mosquito-borne virus estimated to infect 100 million people each year [16, 17]. Four viral serotypes (DENV1–4) exist. Primary infection with a specific serotype confers long-term immunity against subsequent infections of that same serotype, and there is strong empirical support for short term, temporary protection against other serotypes [18–21]. Additionally, other mechanisms such as antibody-dependent enhancement also potentially mediate the dynamics of incidence [22–24].

Multiannual patterns in dengue observed in any location arise as the result of a complex interplay of various factors, including: climate, through its effect on the vector and transmission efficiency; predator-prey dynamics between the virus and the host; the interactions between different serotypes and strains of dengue; spatial patterns in host structure, dynamics, and movement; and viral factors [25–28]. In ascribing drivers to interannual patterns in dengue dynamics, studies have often viewed extrinsic factors, such as climate, and intrinsic factors, particularly the dynamics of immunity, as competing alternative hypotheses [26, 29–35]. However, immunity clearly provides a negative feedback where increases in transmission due to favorable climatic conditions can lead to decreases in transmission in future time periods through the protective effects of immunity, leading to multiannual dynamics in many systems [36, 37].

Many of the same factors involved in generating multiannual dynamics can also produce synchrony across locations [6]. Again, both intrinsic (specifically host movement between locations), and extrinsic (environmental; the “Moran effect”) have the potential to not only synchronise dynamics across space, but also produce variation in the degree of synchrony over time [13–15]. Empirical studies have tended to focus on climate [34, 35]. Van Panhuis *et al*. [34], for instance, detected synchronous dengue outbreaks across Southeast Asia in 1997–1998, a period of elevated temperatures and a strong El Niño event.

The environment, and especially temperature, is *a priori* expected to be significant in shaping the dynamics of dengue [38–40]. Through its effect on metabolic rates, temperature strongly influences many mosquito life history traits (e.g., biting rates, population growth rates, mortality rates; [28, 39]), and therefore, their population dynamics [41]. Projections suggest that increasing temperatures may expand the range in which dengue is efficiently transmitted [42–44]. However, the impact of changing temperature regimes on dengue in locations where the pathogen is already endemic is less well studied. It is in these areas which, even in projections to 2050 or 2080, comprise a majority of the world’s population at risk of dengue, that the impacts of climate change might be greatest [42–44].

Here, we examine the spatio-temporal dynamics of dengue in Thailand between 1968 and 2018. We characterise multiannual cycles and uncover an interesting spatial phenomenon in which Thailand has experienced periodic synchronizations of dengue incidence. In contrast to previous studies, which tended to focus on either extrinsic or intrinsic drivers to explain observed patterns, we hypothesise that immunity constitutes a strong dynamical filter necessary to understand impacts of temperature on dengue. For this reason, we adapt a mechanistic, temperature-dependent, four-serotype dengue model to disentangle how temperature and dynamics of immunity interact to generate periodic synchronizations. We focus on temporary cross-protection between serotypes because of the clear empirical support, although there are other mechanisms, such as antibody-dependent enhancement, that could conceivably also play a role.

## Results

### Empirical patterns in multiannual cycles and synchrony

#### Multiannual cycles in dengue

Fig 1a shows a heatmap of the monthly number of dengue cases by province over the 51 years, in which seasonal outbreaks are clear, but in which some years (e.g., 1998–1999 or 2001) had larger outbreaks throughout the country than other years. It is these latter multiannual patterns that we are interested in here. Specifically, we focus on cyclical patterns with timescales between 1.5 and 5 years (i.e., patterns in which years with larger than average outbreaks are separated by 1.5 to 5 years) because this range encompasses the timescales that have been previously reported for dengue dynamics (see Materials and Methods for a more detailed rationale). To characterise multiannual cycles, we applied continuous wavelet transforms (WTs). WTs quantify the importance of different timescales in a time series over time, and with them, time series can be reconstructed only using only the multiannual components of observed patterns. Reconstructions of the dengue time series confirm the presence of distinct multiannual patterns (Fig 1b). We quantify which multiannual timescale is most important in characterising dengue dynamics in each province and for each point in time (the “dominant timescale”), revealing both that the dominant multiannual timescale tends to change over time (varying between 1.5 and 4.5 years), and that at any given point in time, the dominant timescales can differ substantially across provinces (Fig S6b in Supplementary information (SI)). The presence of multiannual timescales has been previously observed [27, 34], but here, with the benefit of longer time series, we show that these change over time and space.

**Fig 1.**
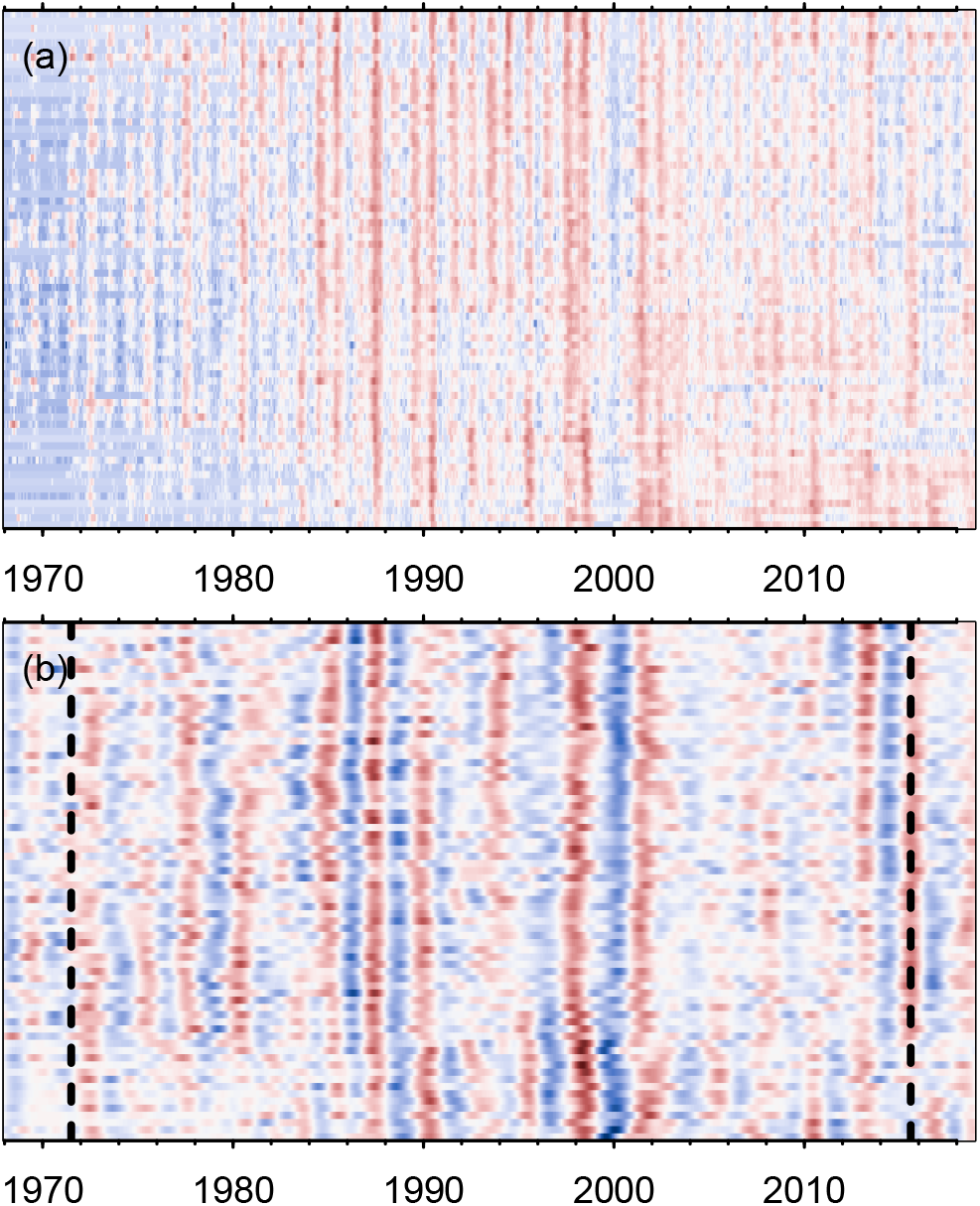
Dengue multiannual cycles. Heatmaps of (a) ln number of cases and (b) reconstructions of time series using multiannual components only, per province arranged from north (top) to south (bottom). To improve clarity, values for each province were normalised to standard scores (*µ* = 0,*σ* = 1) in all panels. Blues (respectively reds) are lower (respectively higher) numbers, and whites correspond to a standard score of zero (the mean for each province). Edge effects in the wavelet transforms may influence results before and after the vertical dashed lines in (b) (see Materials and Methods).

#### Synchrony in dengue

We estimate spatial synchrony in dengue cases using a range of methods; we here focus on one approach, wavelet mean fields (WMFs; [45, 46]), but provide details and results for the others in Section “Perspectives on synchrony” in SI. WMFs measure synchrony as a function of both timescale and time; they indicate timescales and time points at which both phases and magnitude of oscillations are consistent (or more synchronous) across provinces. The WMF for dengue haemorrhagic fever cases across Thailand describes a system that appears to fluctuate in and out of synchrony (Fig 2b), results that are consistent with the travelling waves observed across Thailand [27] and Southeast Asia [34]. The synchrony in dengue is statistically significant at all times, meaning that even when synchrony is lower (the whiter areas in Fig 2b), the degree of synchrony is still greater than would be expected in an asynchronous system.

**Fig 2.**
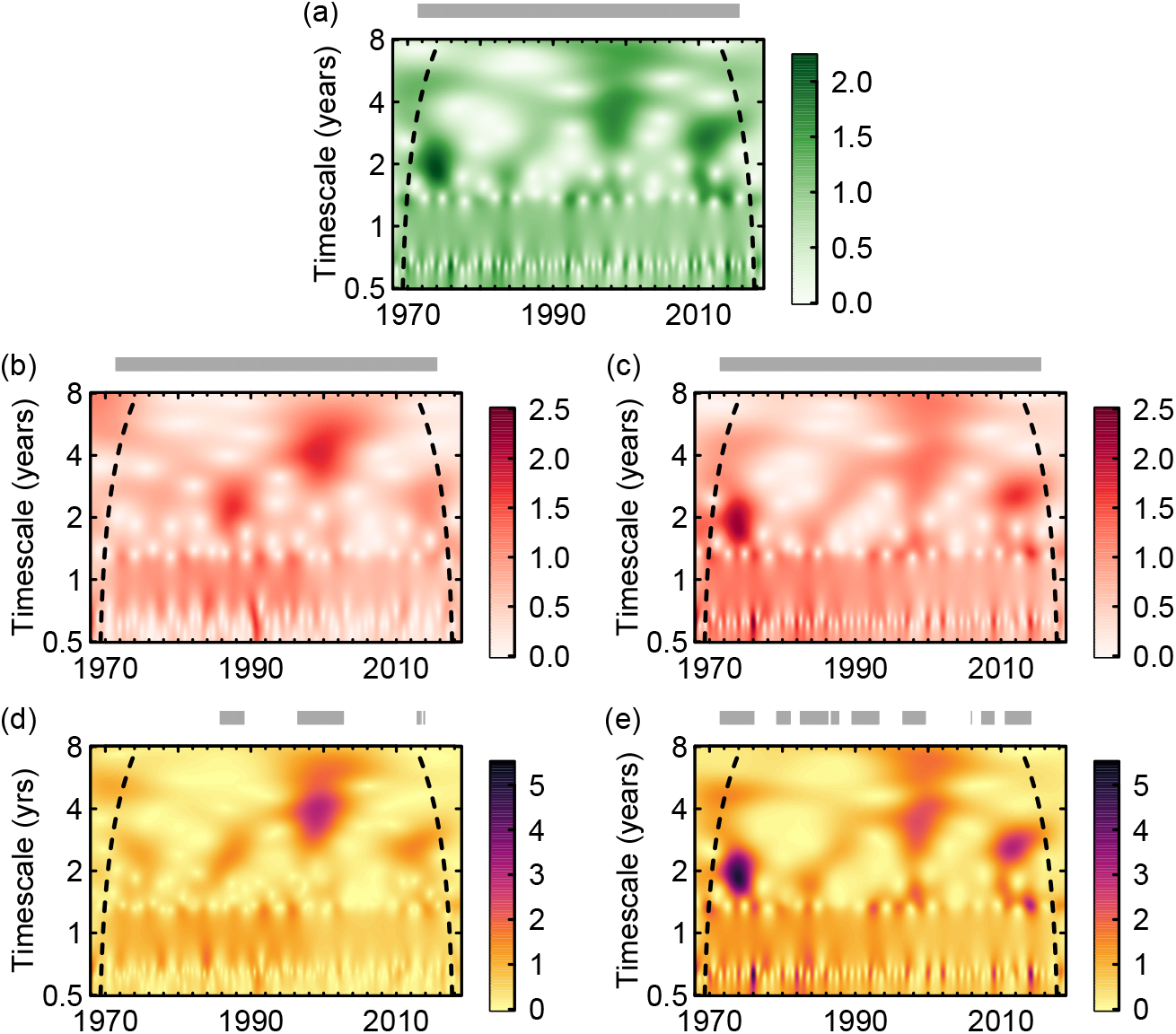
(Cross-)wavelet mean fields. Wavelet mean fields for (a) temperature, and ln-transformed dengue cases from the (b) passive surveillance data and (c) model output. Cross-wavelet mean fields between (d) temperature and ln-transformed dengue cases (data) and (e) temperature and ln-transformed model output. Model output assumes a mean cross-protection of one year (see Fig S13 in SI for results using other mean cross-protections). The same temperature time series are used for both (d,e). Grey bars above panels indicate the times for which phases (in a–c) and phase difference angles (in d,e) are highly consistent across provinces and statistically significant (see Material and Methods). In (a–c), higher values in the mean fields indicate timescales and points in time where the phases are more consistent across provinces, and where the amplitudes of oscillations are more correlated. In (d,e), higher values correspond to timescales and points in time where the agreement between dengue and temperature is itself more consistent across provinces. Fig S14 in SI shows which multiannual timescales dominate the (C)WMFs for each panel in this Figure. Edge effects in the WTs may influence results before and after the dashed lines (see Materials and Methods).

Furthermore, moments of greater synchrony take place at different timescales. For example, while the synchrony event of the late 1980s occurs with a timescale of *≈* 2.2 years, the one taking place around the year 2000 has a timescale of *≈* 4.1 years. These results are further corroborated using four additional approaches to estimate synchrony (Fig 3); all methods describe a system that appears to fluctuate in and out of synchrony (Section “Perspectives on synchrony” in SI). In all cases, periods of greater synchrony appear to coincide with larger outbreaks (e.g., comparing Fig S6a and c in SI), as also previously observed [34, 47].

**Fig 3.**
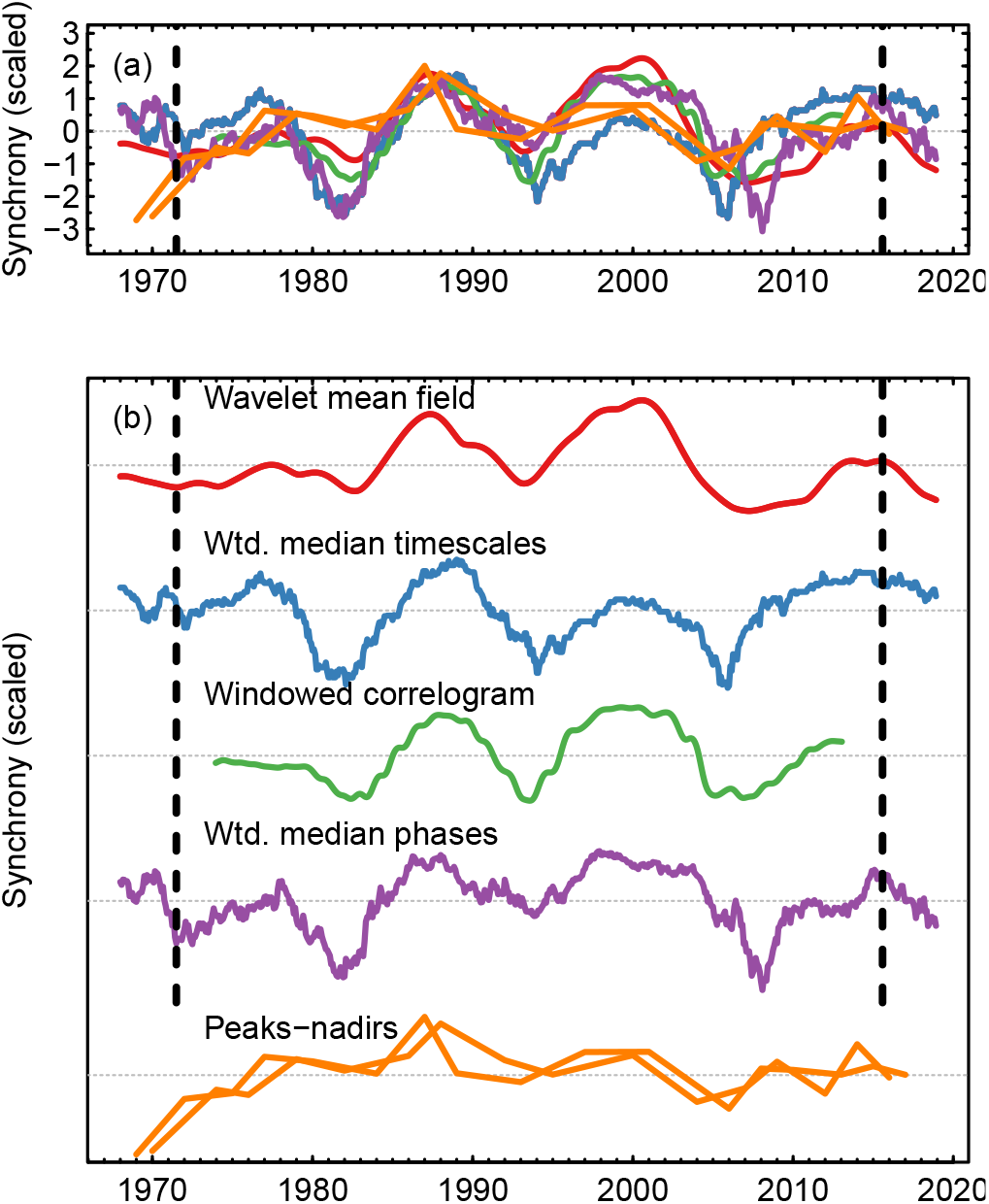
Comparison of all measures of synchrony in dengue. For details on these measures of synchrony, see Section “Perspectives on synchrony” in SI. The curve for wavelet mean fields was obtained by taking the mean WMF of Fig 2b across multiannual timescales, for each point in time. To facilitate comparisons, all metrics were normalised to standard scores, and where necessary, time series were flipped along the y-axis so that higher values always equate to greater synchrony. Note that for the metric taken from the distribution of distances of peaks and nadirs, there are two time series in the same colour (one for peaks and another for nadirs). The two panels show the same time series, but (b) separates them for clarity. Edge effects in the WTs may influence results before and after the vertical dashed lines.

#### Synchrony in temperature

We calculated the temperature WMF across provinces for the same time period as the dengue passive surveillance data (1968–2018). As expected, synchrony in temperature was significant at all points in time. The WMF for temperature describes a system similar to that found in dengue, where temperature fluctuates in and out of synchrony at multiannual timescales (Fig 2a; for results using alternative approaches see Figs S10 and S11 in SI). Furthermore, the moments of greater synchrony appear to align, both in timing and timescale, with those of dengue (Fig 2a,b).

If the patterns in synchrony in dengue were to be directly linked to patterns in synchrony in temperature, we would expect the magnitudes of the two respective WMFs (in the multiannual time scales, and for all points in time) to correlate positively and significantly. We found a Pearson correlation of 0.21 (P = 0.060; see Material and Methods for significance testing) and a Spearman correlation of 0.20 (P = 0.069). Thus, while correlations were positive, they were not statistically significant. However, see the result for WMFs of dengue data and a mechanistic model based on temperature in ‘Driving the model with actual temperature time series’ below.

To address whether the patterns of synchrony in temperature were consistent with those observed in the dengue cases in greater detail, we extended the methods of Sheppard *et al*. [45, 46] to calculate cross-wavelet mean fields (CWMFs). Cross-wavelets quantify the similarity in two time series’ (temperature and dengue cases) wavelet power and phase angles at each timescale and time point, so the CWMFs (the weighted mean cross-wavelet across provinces) quantify the consistency of this similarity across locations. Results in Fig 2d (in which darker colours mean greater consistency between dengue and temperature across locations) suggest that temperature plays a crucial role particularly when synchrony in dengue was high (as demonstrated by the statistically significant consistency in phase angles between dengue and temperature during synchrony events).

### Periodicity of synchronization events in dengue and temperature

Our analyses describe a system that oscillated in and out of synchrony (Figs 2b, 3). To describe the periodicity of the degree of synchrony itself, we collapsed WMFs to a time series by taking the mean WMF across multiannual timescales for each point in time (red line in Figure 3b). We then normalised the resulting time series to standard scores, and applied WTs.

Because these time series of synchrony contain, at most, four distinct cycles (Fig 3), we chose to characterise their periodicity by averaging the wavelet power over time, by taking the mean wavelet power per timescale. The periodicities that dominated the time series of synchrony of dengue and temperature in Thailand, where synchrony was quantified using WMFs, were 12.6 and 12.5 years, respectively (e.g., the main periodicity of the cycles shown in the red line in Fig 3 is 12.6 years). As a point of comparison, the periodicity of synchrony in dengue, where synchrony is estimated using alternative methods such as weighted median timescales or windowed spline correlograms (Section “Perspectives on synchrony” in SI) was 12.0 years in both cases.

### Simulations to disentangle the roles played by temperature and immunity

#### Mechanistic dengue model and simulations

While our results may suggest a relatively straightforward statistical relationship between dengue cases and temperature (Figs 2 and S30 in SI), we wanted to explore the mechanisms by which temperature could generate asynchronous and synchronous dynamics, while accounting for intrinsic factors, such as the interaction between serotypes. Immunity is central in shaping dengue dynamics; temporary cross-protection between serotypes alone can give rise to a qualitatively wide range of dengue dynamics. Any effects of temperature on dynamics must therefore necessarily be viewed through the lens of the dynamics of immunity. To this end, we used a simple, temperature-dependent four-serotype differential equation dengue model (see Materials and Methods). The model, building on that of Huber *et al*. [41], explicitly encodes the temperature dependence of vector traits, which in turn allows for temperature to drive transmission and dengue dynamics, while at the same time allowing for simple, cross-protective interactions between serotypes of varying mean lengths of time.

The patterns quantified so far describe a system that fluctuates in and out of synchrony. When seeking potential drivers for the observed patterns, these would need to account for both asynchrony, and thus the ability to produce a range of spatially heterogeneous dynamics, and synchrony, during which dynamics are more homogeneous.

We ran three sets of simulations to see whether we could better understand how temperature might drive patterns in synchrony, using both real temperature time series and simplified experiments (Table 1; also see SI). First, in simulation 1 we used the Global Historical Climatology Network version 2 and the Climate Anomaly Monitoring System (GHCN CAMS) temperature time series for each province as input for the model, with the objective of reproducing observed spatio-temporal features (specifically the synchrony events; simulation 1). Next, for simulations 2 and 3, we wanted to delve into the specific mechanisms through which temperature may produce asynchronous and synchronous dynamics. The hypothesis that temperature can produce asynchronous dynamics relies on different temperature (and thus transmission) regimes having the potential to generate diverse enough multiannual oscillations across locations. To this end, we devised simplified experiments across five idealised hypothetical locations along a transect in Thailand, going from high mean temperatures and low seasonal variability, to low mean temperatures and high seasonal variability (see Figs S34–S36 in SI). For simulation 2, we used seasonal sine curves as temperature inputs (with no multiannual cycles), and ran simulations for the five locations. Finally, for simulation 3, we built on simulation 2 by introducing a single multiannual fluctuation in the temperature time series, at the same time across the five locations, superimposed over the seasonal cycles. The multiannual fluctuation had varying timescales (2–5 years), and the amplitude of the multiannual wave was a proportion (10%–40%) of the seasonal amplitude. The objective was to determine whether the common multiannual fluctuation across locations would be sufficient to produce synchrony in an otherwise asynchronous system (simulation 3).

**Table 1.** Simulations used in this study. We use the mechanistic dengue model, with different temperature inputs and numbers of locations *N*, to address specific but related questions (column Aim). Each location was run as an independent simulation (no host movement between locations), using the same host birth and death rates and population sizes and starting conditions, such that the only difference across locations was temperature (see section “Further details on simulation studies” in SI). In all cases, simulations were run assuming mean cross-protections between dengue serotypes of six months, one year, and two years. Outputs had a monthly resolution.

| Sim. | $N$ | Temperature input | Aim |
| --- | --- | --- | --- |
| 1 | 72 | Real temperature time series | Reproduce empirical patterns using temperature as the input. |
| 2 | 5 | Seasonal sinusoids | Reproduce asynchrony using different thermal regimes alone. |
| 3 | 5 | Seasonal sinusoids, with a superimposed single multiannual fluctuation (with a timescale of 2–5 years). | Reproduce synchrony in an asynchronous system and explain contributions of intrinsic and extrinsic factors. |

### Driving the model with actual temperature time series

The mechanistic model, driven by temperature with the mediation of temporary serotype cross-protection, was capable of producing qualitatively similar dynamics to those observed in the data (e.g., see Fig S12 in SI), thereby supporting the hypothesis that temperature plays an important role. For simulation 1 (using temperature time series for the 72 provinces), the WMF for the model output had statistically significant synchrony at all times. The Pearson and Spearman correlations between the model output and the data WMFs (Fig 2c and b) were 0.29 (P = 0.023) and 0.32 (P = 0.016), respectively. These correlations show a statistically significant association between the overall patterns in synchrony in dengue cases and model output, supporting the hypothesis that temperature is likely a strong driver of synchrony in dengue. For different assumptions on the duration of cross-protections, the correlation coefficients between the model output and data WMFs were marginally lower, and P values were higher (Table S1 in SI). The phase angles between temperature and model output across provinces were, as expected, consistent and statistically significant over most time points (Fig 2e). Both the WMF and CWMF for the model output contain a notable synchrony event in the early-1970s, reproducing that found in temperature; this feature is missing from the WMF and CWMF estimated using the data (Fig 2b,d). WMFs and CWMFs for model outputs using different assumptions on cross-protection are shown in Fig S13 in SI, and show that with longer mean cross-protections (e.g., two years), the degree of synchrony in the model output, and consistency in the phase angles between modelled dengue and temperature, is lower than with shorter cross-protections.

### A temperature gradient can generate asynchronous dengue dynamics

For the idealised experiments of simulation 2, we found that with at least a mean of one year temporary cross-protection between serotypes, the resulting dengue dynamics include multiannual cycles (Fig 4a–c), cycles that are completely attributable to the dynamics of immunity given that here there are no multiannual cycles in temperature. The periodicity of these cycles (and their amplitude), however, changed as a result of the temperature regime. As temperature conditions are optimised for transmission (i.e., as mean temperatures increase, approaching the higher temperatures that characterise temperature in, for instance, Bangkok; Fig S36 in SI), the periodicity of multiannual components in dengue go down (i.e., multiannual cycles are more frequent), and the power at the multiannual timescales increases. At the highest mean temperature, with a mean of two years of cross-protection, seasonal cycles all but disappear (Fig 4c) due to two main trends. On the one hand, as cross-protection becomes longer, the power of the multiannual cycles (and therefore their amplitude) increases (Fig 4a–c). On the other, the seasonal variation in transmission is most limited at the highest mean temperatures (Fig S37 in SI), thus producing more limited seasonal variation in dengue cases. Taken together, these two trends mean that dynamics become increasingly dominated by the multiannual cycles produced through cross-protection, and less so by seasonal variation in temperature. While the periodicity of the multiannual cycles for each temperature regime changes little between means of one and two years of cross-protection, their prominence does: with longer cross-protection, these cycles clearly increase in mean wavelet power. The range of dynamics produced across locations under all assumptions of cross-protection are sufficiently diverse: no synchrony is detected (e.g., Fig 4d).

**Fig 4.**
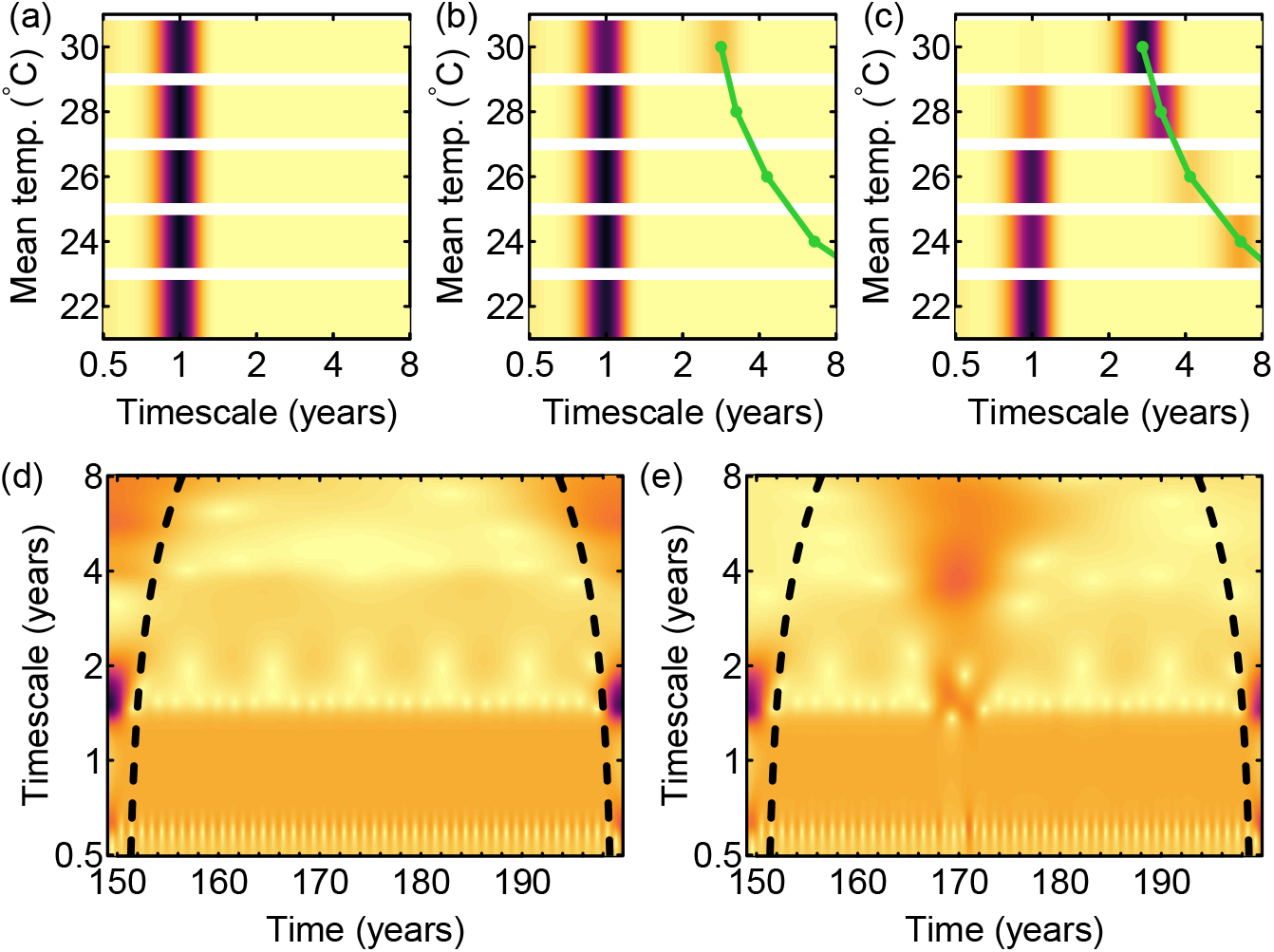
Multiannual periodicities and synchrony for models driven by simple seasonal sine curves. (a–c) Mean wavelet power for simulation 2 experiments (seasonal temperature input with no multiannual components; Table 1) per timescale (over time) for each of the five hypothetical locations (rows within each panel), assuming a mean cross-protection of (a) six months, (b) one year, and (c) two years. (d) Mean wavelet field for the simulations in (b) (i.e., assuming a cross-protection of one year), and (e) mean wavelet field for the same simulation, except for the introduction of a single four-year fluctuation across all locations on year 168. The amplitude of the multiannual fluctuation here is 20% of that of the seasonal cycle. Darker colours indicate greater wavelet power (colours in a–c and d–e are (separately) on the same scale). Green lines indicate peaks in mean wavelet power over multiannual timescales. Edge effects in the WTs may influence results before and after the dashed lines in (d,e). See Figs S15–S28 in SI for more complete results (across all cross-protections, and timescales and amplitudes of the multiannual fluctuation).

### Common multiannual fluctuations in temperature can synchronise dengue dynamics

Because different temperature regimes produce distinct dengue dynamics (Fig 4a–c), one simple explanation for synchrony could be that temperatures across the country converge to a more similar temperature regime during those periods, resulting in more synchronous dengue dynamics. However, this has not been the case in Thailand (Fig S32 in SI). The next hypothesis is that a common multiannual fluctuation in temperature can synchronise dengue dynamics (simulation 3). The result for simulation 3 was that when the amplitude of the multiannual fluctuation in temperature was at least 20% of the seasonal cycle, the five locations were temporarily synchronised (Fig 4e), under all assumptions on cross-protection. Synchrony in dengue occurred at approximately the same timescale as that of the multiannual fluctuation in temperature, so, for example, when the multiannual fluctuation in temperature was a single four-year cycle, synchrony in dengue was detected at a four-year timescale. However, when the multiannual fluctuation had an amplitude 10% that of the seasonal cycle and mean cross-protection was two years, synchrony was not detectable (Fig S28 in SI).

### The roles played by immunity and temperature

The synchronisation achieved in the simulation 3 experiments varied as a function of the mean duration of cross-protection: the degree of synchrony fell with a longer cross-protection (Figs S20–S28 in SI). To explain why, we need to better understand how the multiannual timescales produced by intrinsic and extrinsic factors interact. Comparing simulations with and without a single multiannual fluctuation in temperature allows us to quantify how the multiannual timescales in dengue change specifically as a result of the multiannual fluctuation in temperature. We found that while the periodicity of the multiannual fluctuation in temperature is generally detectable in the resulting dengue dynamics, its importance relative to the timescales of the intrinsic dynamics (i.e., the dengue multiannual timescales attributable to cross-protection alone) vary as a function of both mean temperature and mean cross-protection (Fig 5). Dengue appears to be more insensitive to multiannual fluctuations in temperature with increasing mean temperatures and cross-protections. Specifically, as the duration of cross-protection and/or mean temperatures increase (and as seasonality in temperature is reduced), the power of the multiannual intrinsic dynamics increases, such that the effect of the multiannual fluctuation in temperature is overwhelmed. In our simulations, this is particularly the case for mean temperatures of 30°C and a mean cross-protection of two years (Fig 5).

**Fig 5.**
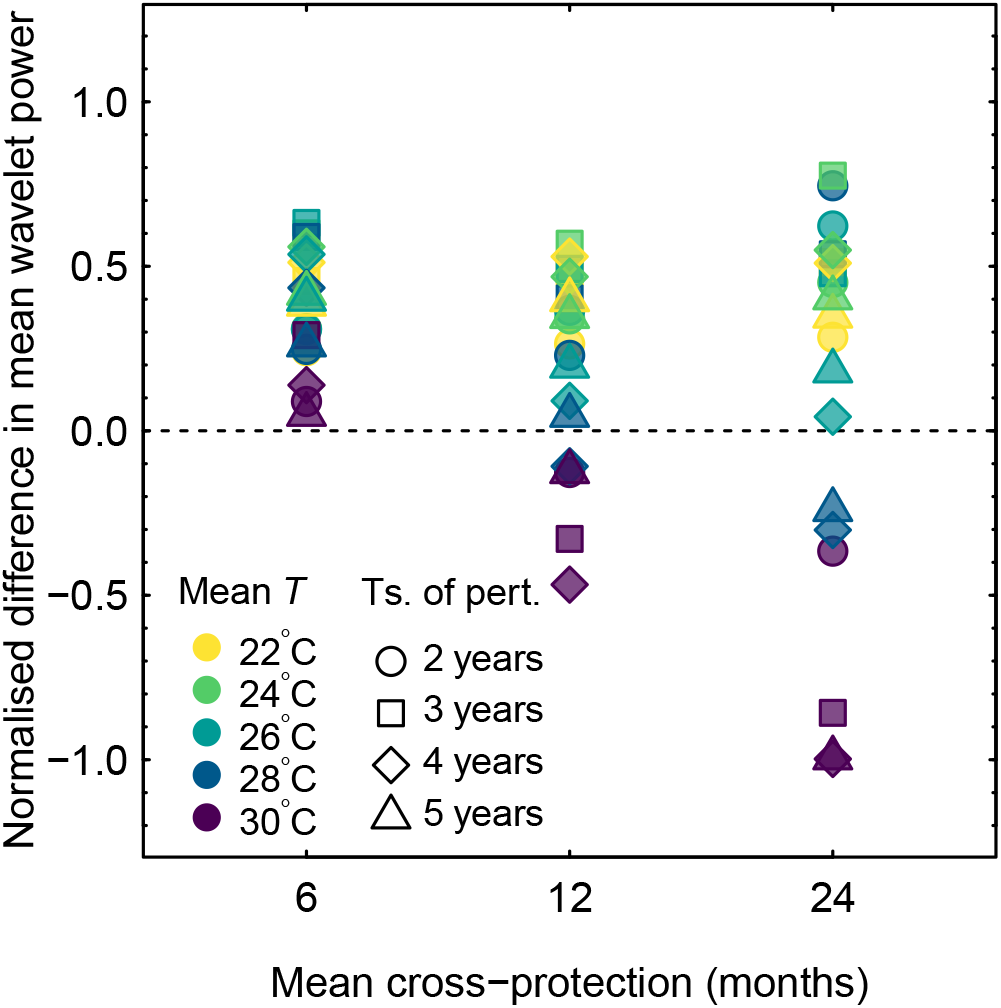
Comparing multiannual timescales in dengue caused by cross-protection and by multiannual fluctuations in the environment using simulations. We compare the mean wavelet power of dengue at the timescale of the environmental multiannual fluctuation, with the mean wavelet power of dengue at the timescale at which wavelet power is maximised without a multiannual fluctuation in temperature (the multiannual timescale attributable to cross-protection alone). Positive values mean that the multiannual fluctuation in temperature plays a more important role driving dengue multiannual dynamics, and negative values mean that dengue dynamics are less sensitive to the multiannual fluctuation in temperature. For example, for a mean temperature of 30°C, a multiannual fluctuation with a timescale of four years, and a mean cross-protection of two years, we compared the mean wavelet power at a timescale of four years (that of the multiannual fluctuation), with the mean wavelet power at a timescale of 2.7 years (the multiannual timescale at which wavelet power was maximised with no multiannual fluctuation in temperature, i.e., the peak on the top row of Fig 4c). In this example, the mean wavelet power was distinctly greater at the 2.7 year timescale than at the 4 year timescale, meaning dengue dynamics were insensitive to the multiannual fluctuation in temperature. Differences in wavelet power shown here have been normalised against the maximum value for each timescale of the multiannual fluctuation, to improve clarity. Overall, longer cross-protection (the right column of points) and mean temperatures (purple colors) are more likely to lead to a system dominated by intrinsic dynamics, and less sensitive to multiannual fluctuations in temperature. Mean wavelet power is estimated for the duration of the multiannual fluctuation only (so for a four-year multiannual fluctuation in temperature, mean wavelet power is estimated during those four years only). Here, the multiannual fluctuation has an amplitude 0.2 times that of the seasonal cycle, but the patterns are qualitatively similar for the other amplitudes of the multiannual fluctuations.

It is also worth noting that the multiannual fluctuation in temperature is not necessarily straightforwardly reproduced in dengue dynamics. Instead, it elicits a transient response in dengue cases that can extend in time beyond the multiannual fluctuation in temperature, as can be clearly seen when plotting the phase angle between simulations with a multiannual fluctuation in temperature, and those without (Fig S29 in SI). These show both that the effect of the multiannual fluctuation in temperature can be lasting, and also that the effect of the multiannual fluctuation is reduced with longer cross-protections and higher mean temperatures.

This interaction between temperature and cross-protection explains why with longer cross-protection, the degree of synchrony appears to go down: as cross-protection becomes longer, locations with higher mean temperatures are more likely to be dominated by the intrinsic timescales, and as a result, the ability of a multiannual fluctuation in temperature to synchronise locations is reduced.

## Discussion

Using multiple approaches, we have described an interesting spatial phenomenon in dengue: the country has repeatedly and periodically moved in and out of synchrony in dengue cases. In general, periods of greater synchrony coincided with larger outbreaks. When analysing temperature time series across Thailand, we found a similar pattern in spatial synchrony. Although the correlation between the overall patterns in synchrony in temperature and dengue cases was not statistically significant, the relationship between temperature and dengue cases was particularly consistent across provinces during times of greater synchrony. Accounting for the dynamics of immunity is essential for understanding dengue dynamics; any effects of temperature on dynamics are modulated by immunity. To gain a more mechanistic understanding of how temperature interacts with the dynamics of immunity to produce the observed patterns in synchrony, we adapted a mechanistic dengue model with temporary cross-protection. When running the model using the observed temperatures across Thai provinces, the patterns in synchrony in the resultant model output dengue time series correlated positively and significantly with observed patterns in dengue cases from the passive surveillance data. The fact that this correlation was statistically significant, but the one between the patterns in synchrony in dengue cases and temperature was not, further highlights the importance of nonlinearities in the way temperature affects dengue and dynamics of immunity. While (a sufficiently long) cross-protection alone can produce multiannual cycles, we found that their periodicity was modulated by temperature; different temperature regimes produced characteristically different dengue dynamics. However, although the timescales of the multiannual periodicities of dengue varied little, their power became more prominent with longer cross-protections. This result sheds some light on the roles played by both temperature and dynamics of immunity, and suggest that both are essential to understand patterns in dengue dynamics. We also found that a range of temperature regimes produced asynchronous dengue dynamics, but the introduction of common multiannual fluctuations across a range of different temperature regimes was sufficient to synchronise the otherwise asynchronous system. The degree of synchrony depended on the duration of cross-protection: as the duration of cross-protection increased, the multiannual cycles produced by cross-protection became more prominent compared to those produced by multiannual fluctuations in temperature, particularly at higher mean temperatures, resulting in a lower degree of synchrony across locations. With longer cross-protections, a larger proportion of the population is temporarily protected and not susceptible to infections, and thus unaffected by variations in transmission. However, with less than two years of mean cross-protection, synchrony in temperature can synchronise an otherwise asynchronous system.

Our results highlight how ongoing climate change, together with other concurrent changes, may be affecting dynamics in areas where the disease is endemic and a majority of the population at risk, now and in future projections, lives.

Most locations’ temperature appear to be increasing in mean and decreasing in seasonal variation (Fig S33 and S36 in SI). If these trends were to be maintained, our results suggest that dengue dynamics could slowly shift too, and approach a regime characterised by higher average incidence (Figs S15–S17 in SI), and dynamics increasingly dominated by multiannual, rather than seasonal, cycles. Multiannual cycles may also be, to greater extents, dictated by intrinsic factors over the multiannual timescales present in temperature. These trends may also suggest that in the future, synchrony may be driven by increasing similarity in temperatures across Thailand, and less so by synchronous multiannual fluctuations in temperature. However, these hypotheses are contingent on a more precise understanding of the interactions between serotypes and contributions from other intrinsic factors (such as changes in demography and movement of hosts between locations).

There are several caveats to our analyses. We chose to focus on temperature and its interaction with the dynamics of immunity. However, we could not ascribe every pattern in synchrony in dengue to synchrony in temperature. For example, Thailand experienced a strongly synchronous two-year fluctuation in temperature around 1972, and while this fluctuation was reproduced in our model output, it was conspicuously absent from the observed dengue dynamics. There are multiple reasons that could explain this discrepancy. During the earlier part of the records, dengue cases might have been detected less effectively, thus potentially obscuring multiannual patterns. On the other hand, other environmental variables (notably precipitation) and their interaction with temperature are likely important for the dynamics of dengue. For instance, the extent to which a synchronous event in temperature can synchronise multiannual cycles in dengue might depend on the amount of precipitation or timing of the rainy seasons during those years. Additionally, other non-environmental mechanisms might also influence synchrony in dengue. Complex multiannual cycles, synchrony, and variations in the degree of synchrony over time can in theory be produced via, for example, the movement of hosts between locations (weakly-coupled oscillators; [15]). There are also a range of concurrently ongoing changes in Thailand that may also influence multiannual dynamics and synchrony. For instance, the birth rates across the country have been declining [48], and the rates of decline may be spatially heterogeneous. Similarly, long-term trends in population densities across the country could also contribute to the observed spatio-temporal patterns. Exploring how these factors may interact with temperature and contribute to the observed spatio-temporal patterns deserves further attention. Finally, the mechanistic model we use here necessarily makes simplifying assumptions (see “Temperature-dependent dengue model” in SI). Further work is required to understand which of these assumptions may significantly affect the dynamics produced.

## Materials and methods

### Data

We use data provided by the Thai Ministry of Public Health in their Annual Epidemiological Surveillance Reports. Monthly dengue case counts are given per province, starting in January 1968. Prior to 2003, the counts provided combine the cases of dengue fever, dengue shock syndrome, and dengue haemorrhagic fever (DF, DSS, and DHF, respectively), while separate counts are given for each category thereafter. For 2003 onwards we use DHF cases only given these are the cases that are most likely to lead to severe outcomes and that these are the most likely to have been reported prior to 2003, although results change little when including DF and DSS. Starting in 1982, five new provinces were created. To maintain consistency across the time series, cases for these new provinces were added back to the provinces they were created from, keeping the total number of provinces (72) constant over time.

Temperature time series were obtained from the GHCN CAMS gridded (0.5° by 0.5° resolution) monthly mean temperature data set (provided by NOAA/OAR/ESRL PSD, Boulder, Colorado, USA, from https://www.esrl.noaa.gov/psd/data/gridded/data.ghcncams.html, downloaded on 18/06/2019). This dataset combines station observations from the Global Historical Climatology Network v2 and the Climate Anomaly Monitoring System [49] (Figs S31 and S32 in SI). The time series start in 1948. We downloaded shapefiles for Thai provinces from https://data.humdata.org/dataset/thailand-administrative-boundaries on 12/02/2019, and estimated province centroids using function ‘calcCentroid’ in R package ‘PBSmapping’ v2.72.1. A time series for each province was then obtained by using the grid point nearest to the centroid of each province.

### Wavelet transforms

We applied continuous wavelet transforms (WTs) to explore how the oscillatory behaviour of dengue cases has changed over time. WTs have now been applied extensively in the study of infectious disease dynamics [31, 34, 50–52] and more broadly in ecology [45, 46]. WTs decompose time series into frequency components, but, as opposed to Fourier transforms, WTs are also localised in time. Thus, WTs can be used to characterise non-stationary time series and how the relative importance of different frequencies change over time. The basis for the WT is a “mother” wavelet, a wave localised in both frequency and time (i.e., of limited duration), which is then scaled or stretched (for the frequency component) and shifted along the time axis (for the temporal component) to derive a set of “daughter” wavelets. The WT can then be understood as a correlation between (or more specifically the convolution of) the time series and the set of daughter wavelets [53].

We use the WT as implemented in R package ‘WaveletComp’ v1.1 – details are provided in the package documentation [54] and code, and a summary is given here. The package uses a Morlet mother wavelet, *ψ*(*t*) = *π*^*−*1*/*4^ exp(*i ω t*) exp(−*t*^2^*/* 2), where *t* is time (*t* = 1, …, *T*) and *ω* is the angular frequency, set to 6 radians *t*^*−*1^ [54]. The WT, *W*, at a point in time *τ* and scale *s* (proportional to period, the inverse of frequency) is:

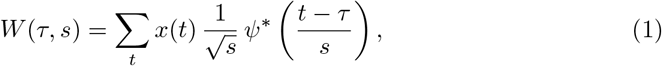

where *x*(*t*) is the original time series, and * denotes the complex conjugate. The modulus of *W* (*τ, s*), |*W* (*τ, s*) |, gives the local amplitude *A* at time point *τ* and scale *s*. For any plots of the WT or calculations performed directly on it, we correct *A* so that *A*(*τ, s*) = *s*^*−*1*/*2^ |*W* (*τ, s*) | [55]. *A*(*τ, s*)^2^ is the wavelet energy density. *W* (*τ, s*) also yields the instantaneous local wavelet phase. Although the WT contains significant redundancy in time and scale, it is possible to reconstruct the original time series (or time series containing specific scales only) on the basis of summing over the real part of the wavelet transform [53].

For these analyses, prior to performing the WT, a value of one was added to time series of dengue cases prior to ln-transforming and normalising to standard scores (mean = 0, standard deviation = 1). Time series of dengue cases were not detrended prior to calculating wavelet power for two reasons: detrending a time series with zero values produced artificially odd patterns. Time series of temperature were not transformed; long-term nonlinear trends were removed by taking the residuals of a local polynomial regression (using function ‘loess’ with a span of 0.75), and the detrended time series were subsequently normalised, as above. Due to the finite length of the time series, whenever the wavelets extend beyond the edges of the time series, estimates of transform coefficients become less accurate. This issue is exacerbated as scales increase, due to wavelets extending further in time [53, 54]. Regions where these edge effects are present (often referred to as the “cone of influence”) are clearly indicated in wavelet plots by thick dashed lines.

While the dynamics of dengue are seasonal, previous studies have also identified distinct multiannual timescales [27, 29, 31, 34, 52]. We are specifically interested in these multiannual timescales; for this reason we focus on timescales greater than 1.5 years. This lower bound also prevents leakage from the annual signal into the multiannual timescales. As the timescales increase, edge effects increasingly dominate and the length of time we can reliably analyse becomes shorter. Furthermore, previous studies [29, 34] typically identified multiannual components at 2–3 years, albeit using shorter time series. For these reasons we here focus on the 1.5–5 year timescales (henceforth, “multiannual” components or timescales refer to this range).

### Wavelet mean fields as a measure of synchrony

Wavelet mean fields (WMFs), provide a measure of synchrony as a function of both scale and point in time (further details can be found in Sheppard *et al*. [45, 46]; only a summary is provided here). More intuitively, WMFs indicate scales and time points at which both phases and magnitude of oscillations are consistent (or more synchronous) across provinces. WMFs are a mean of the power-normalised WTs of each location *n* across all *N* locations (*n* = 1, …, *N*), where we normalised each WT by

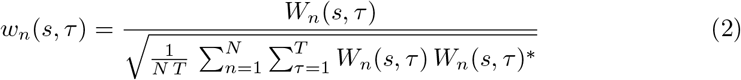

[45, 46].

To test whether synchrony is statistically significant in the multiannual range against a null hypothesis of no synchrony, we focus on the consistency of phases across locations. We do so by estimating the wavelet phasor mean field (WPMF), defined as

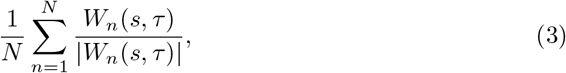

which retains information on the complex phases of the transforms [46]. When different locations have a similar phase, the value WPMF will be large, while when the phase in each location is independent from all other locations, the value of the WPMF will be small. We test significance by generating surrogate datasets where the autocorrelation structure within each time series is preserved. We do this by generating time series with the same Fourier spectrum as the original time series [46]. Following this approach, we produced 1000 surrogate time series, using function ‘surrog’ in R package ‘wsyn’ v1.0.2 [56]. For each surrogate dataset, we produced a WPMF, and then, for each point in time, and across the multiannual scales, we:

1. compared, scale by scale within a single time point, the power of the ‘real’ WPMF with that of all surrogate WPMFs, noted which scales were at least in the 95 percentile, and then calculated the fraction of scales at that point in time for which this was the case (producing a time series of proportions);
2. repeated step 1 but taking each surrogate WPMF in turn, and comparing it to all other surrogate WPMFs (producing a time series of proportions for each surrogate WPMF); and, finally,
3. calculated the points in time for which the time series of step 1 were at least in the 95 percentile compared to the time series in step 2.

## Drivers of synchrony

To assess the association in the patterns in synchrony between the WMFs of dengue cases and temperature, and between the WMFs of dengue cases and modelled dengue output, we perform both Pearson and Spearman correlations on the respective WMF values within the timescales of interest for all points in time (excluding the cone of influence). These correlations quantify the overall similarity in the pairs of WMFs, and when statistically significant, would imply that the two WMFs are related (and thus lend support to the hypothesis that the patterns in synchrony of temperature drive patterns in dengue). For the Pearson correlations, we square-root transform the WMF values. We estimate statistical significance by producing null distributions of correlations. We generate surrogate WMFs for 1000 surrogate datasets of dengue cases, obtained, as above, by preserving the same Fourier spectrum as the original time series of dengue cases, but assuming asynchrony. The Pearson and Spearman correlations between each of these surrogate WMFs and the temperature WMF or the model output WMF produce the null distributions against which we compare the observed correlations.

To determine in greater detail whether the patterns of synchrony in temperature were consistent with those observed in the dengue cases, we extended the method above for WMFs [46] to calculate cross-wavelet mean fields (CWMFs). Cross-wavelets quantify the similarity in two time series’ (temperature and dengue cases) wavelet power at each scale and time point, so the CWMFs quantify the consistency of this similarity across locations. CWMFs are an average of the power-normalised cross-wavelets at each location between temperature and dengue cases. A cross-wavelet *X* between two time series is defined as

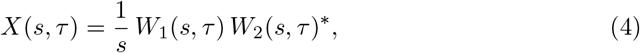

where the cross-wavelet is corrected following Veleda *et al*. [57]. The argument of *X* gives the phase angles between the two time series per time point and scale. For timeseries of temperature and dengue cases with respective power-normalised (eq. (2)) wavelets 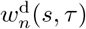 and 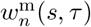, the CWMF is defined as

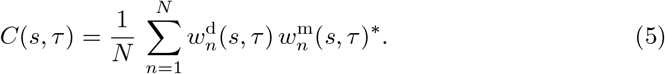

*C*(*s, τ*) will be large if the phase differences between temperature and dengue for each time point and scale are are consistent across locations, and if the amplitudes of the oscillations are correlated. Two unrelated but temporally and spatially autocorrelated variables can quite readily produce patterns in synchrony. For example, if instead of using temperatures for Thailand we were to produce a CWMF using temperatures from a different location, with similar temporal and spatial autocorrelation, we could potentially observe similar patterns in synchrony, despite the two variables being clearly unrelated. For this reason, we test significance in the multiannual timescales using the same three-step approach described above, but here, surrogate dengue datasets are produced that not only preserve the autocorrelation structure of the time series, but also the cross-correlation structure across locations (i.e., synchrony-preserving surrogates). We do this by generating time series with the same Fourier spectrum as the original time series (as above), but then adding the same random uniformly distributed phase at each frequency across all time series [46]. As above, we test significance by focussing on the consistency of phase angles between dengue and temperature. Using these surrogate dengue datasets, cross-wavelets are estimated relative to the real temperature dataset, and these in turn are used to produce the surrogate cross-wavelet phasor mean fields, which we use for comparison.

## Supporting information

This article contains supporting information.

## Acknowledgments

This project was funded by NIH National Institute of Allergy and Infectious Diseases grant R01 AI114703-01. We thank Matt Hitchings and Jessica Metcalf for comments on the manuscript and valuable discussions.

## Disclaimer

Material has been reviewed by the Walter Reed Army Institute of Research. There is no objection to its presentation and/or publication. The opinions or assertions contained herein are the private views of the author, and are not to be construed as official, or as reflecting true views of the Department of the Army or the Department of Defense.

